# Evaluating Augmentation Approaches for Deep Learning-based Major Depressive Disorder Diagnosis with Raw Electroencephalogram Data^*^

**DOI:** 10.1101/2023.12.15.571938

**Authors:** Charles A. Ellis, Robyn L. Miller, Vince D. Calhoun

**Author notes:** Funded by NIH R01MH123610, NIH R01MH118695, and NSF 2112455.

## Abstract

While deep learning methods are increasingly applied in research contexts for neuropsychiatric disorder diagnosis, small dataset size limits their potential for clinical translation. Data augmentation (DA) could address this limitation, but the utility of EEG DA methods remains relatively underexplored in neuropsychiatric disorder diagnosis. In this study, we train a model for major depressive disorder diagnosis. We then evaluate the utility of 6 EEG DA approaches. Importantly, to remove the bias that could be introduced by comparing performance for models trained on larger augmented training sets to models trained on smaller baseline sets, we also introduce a new baseline trained on duplicate training data to better. We lastly examine the effects of the DA approaches upon representations learned by the model with a pair of explainability analyses. We find that while most approaches boost model performance, they do not improve model performance beyond that of simply using a duplicate training set without DA. The exception to this is channel dropout augmentation, which does improve model performance. These findings suggest the importance of comparing EEG DA methods to a baseline with a duplicate training set of equal size to the augmented training set. We also found that some DA methods increased model robustness to frequency (Fourier transform surrogates) and channel (channel dropout) perturbation. While our findings on EEG DA efficacy are restricted to our dataset and model, we hope that future studies on deep learning for small EEG datasets and on new EEG DA methods will find our findings helpful.

## I. Introduction

The application of deep learning methods to raw electroencephalography (EEG) data is becoming increasingly common. However, developing reliable models can be difficult due to the small size of many EEG datasets [1], [2]. One approach that has been frequently used to solve this problem in other domains is data augmentation (DA), wherein synthetic data is generated for use in model training by duplicating and modifying the training data [1]. Multiple DA approaches have been developed for EEG [1], but their comparative efficacy for the diagnosis of neuropsychiatric disorders with multichannel EEG remains relatively unexplored. In this study, we evaluate 6 different EEG DA approaches and compare their effect upon a model trained for major depressive disorder (MDD) diagnosis. We identify which approaches most enhance performance and explore their effects upon features learned by the model via spectral and spatial explainability analyses.

Deep learning is increasingly being applied to raw EEG data. However, relative to more traditional machine learning applications, deep learning approaches require significantly more data to prevent overfitting [1]. Common applications of deep learning with EEG include sleep stage classification, seizure detection, and brain computer interfacing [3], and there are multiple large publicly available datasets for these applications, which makes the development of robust models more feasible [3]. Nevertheless, there is also a growing body of literature focused on using deep learning with EEG for automated neuropsychiatric disorder diagnosis [4]–[6]. The publicly available datasets for these applications tend to be smaller, and collecting new datasets can be very time-consuming and financially intensive [2]. As such, there is a need for the development and evaluation of new approaches that will enable the creation of reliable models for neuropsychiatric disorder diagnosis on small datasets.

Multiple approaches can aid robust model development on smaller datasets. Two approaches are (1) transfer learning and (2) DA. A growing corpus of transfer learning methods focuses on EEG data [2], [6], [7], wherein models are trained for a task on a large dataset and then transferred to a target task with a smaller dataset. While transfer learning can be effective, it is an intensive process, and the development of standard pretrained EEG models like those in computer vision [8] is still in its infancy. DA offers a potentially faster and simpler approach. DA is often applied in computer vision, and multiple DA approaches have been developed for EEG [1]. Their efficacy has been compared in popular tasks like sleep staging and brain computer interfacing [1]. However, their utility for neuropsychiatric disorder diagnosis remains relatively unexplored.

In this study, we evaluate the efficacy of 6 EEG DA approaches for MDD diagnosis. We train a baseline model without DA, train models with duplicate training data to determine whether simply increasing training set size improves performance, and train models with the 6 DA methods. We lastly use spectral and spatial explainability methods to identify the effect of DA upon the features learned by the models.

## II. Methods

We describe our datasets, preprocessing, model development, transfer learning approach, and explainability analyses. To enable reproducibility, code is made publicly available (https://github.com/cae67/DataAugmentationConference).

### A. Datasets and Preprocessing

We used an EEG dataset [9] recorded at resting state with eyes closed from 30 MDDs and 28 HCs, where each recording lasted for 5-10 minutes, used a sampling rate of 256 Hertz, and used a 10-20 format with 64 electrodes. The data has been used in multiple deep learning studies [4]–[6], [10]. Because the dataset is publicly available, no local Institutional Review Board approval was needed for this study. As in previous studies [5], [6], [11], we used 19 channels: Fp1, Fp2, F7, F3, Fz, F4, F8, T3, C3, Cz, C4, T4, T5, P3, Pz, P4, T6, O1, and O2. The data was low pass filtered at 70Hz and notch filtered at 50 Hz. We downsampled the data to 200 Hz and channel-wise z-scored recordings separately. To increase the number of available samples, we separated the data into 25-second samples with a sliding window approach (step size = 2.5 seconds). Our dataset had 2,942 MDD and 2,950 HC samples.

### B. Model Development Approach

In this study, we developed a baseline model with no DA (Model 1.1), developed models with the training duplicated and triplicated data to examine whether simply increasing the training set size was sufficient to increase performance (Model 1.2, M1.3), and trained models with 6 DA approaches that doubled (M2.1-7.1) or tripled (M2.2-7.2) training set size. We then compared model performance via a statistical analysis.

#### a) Model 1.1 (Baseline) Development

We adapted an architecture [12] that has been used for MDD classification in a number of studies [4]–[6] (Figure 1). Similar to [7], we optimized the hyperparameters with the Hyperband algorithm, a Keras-Tuner [13] automated machine learning approach (10 initial epochs per trial, maximum of 40 training epochs). For Hyperband, we optimize the average validation accuracy (ACC) across 25 folds. When performing cross-validation, we used a subject-wise, group shuffle split approach that assigned 70%, 20%, and 10% of the subjects to training, validation, and test sets, respectively. We used a batch size of 128, Adam optimization (best learning rate of 0.001), and early stopping if validation ACC did not increase for 5 subsequent training epochs. After hyperparameter tuning, we retrained the model using checkpoints to select models from the epoch of each fold with the top validation ACC. When evaluating test performance, we calculated ACC, balanced ACC (BACC), sensitivity (SENS), and specificity (SPEC).

**Figure 1.**
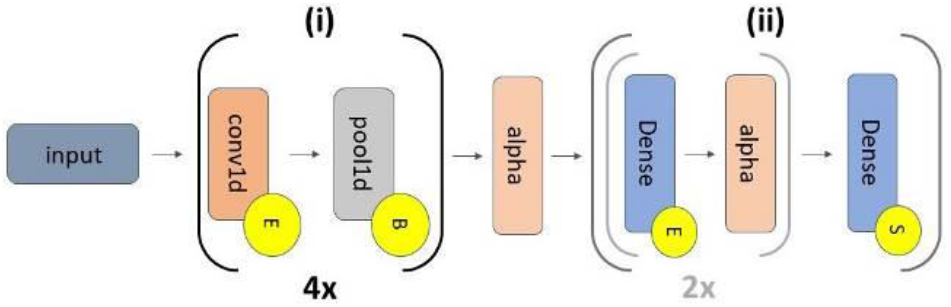
Model Architecture. Models have 2 portions separated by alpha dropout (alpha, rate = 0.4): (i) a feature extractor (repeated 4 times), and (ii) a classifier. The grey subportion of (ii) repeats twice. There are 4 convolutional (conv1d) layers (number of filters = 10, 20, 5, and 20; kernel size = 20) followed by max pooling layers (pool size = 2, stride = 2). Portion (ii) has 3 dense layers (number of nodes = 16, 48, and 2) interleaved with alpha layers (rate = 0.5). Yellow circles with an “E” and “S” or “B” correspond to ELU and softmax activations and batch normalization, respectively. All dense and conv1d layers had max norm kernel constraints (max = 1).

#### b) Model 1.2-1.3 Development

Using the optimal hyperparameters identified for M1.1, we first examined whether simply duplicating (1.2) or triplicating (1.3) the training data in each fold to obtain double or triple the number of training samples was sufficient to improve performance. M1.2 and M1.3 training was virtually identical to that of M1.1, though we did optimize the learning rate of the models.

#### c) Model 2-7 Development

We evaluated 6 DA methods: Gaussian noise (M2), time reverse (M3) [14], smooth time masking (M4) [15], Fourier transform (FT) surrogate (M5) [16], frequency shift (M6) [14], and channel dropout (M7) [17]. When evaluating each approach, we used the hyperparameters identified during M1.1 training and used the cross-validation and other training features of M1.1. However, we also optimized the learning rate of each augmented model with Hyperband and optimized the DA parameters. We doubled or tripled the training set size with each DA approach (i.e., duplicated the training data once or twice, applied DA, and trained on the combined original and augmented data). Models trained with DA and doubled or tripled datasets are designated with a .1 and .2, respectively.

Due to page constraints, we do not go into exhaustive detail on each of the DA approaches. We instead cite the studies that originally introduced or that have used the methods. [1] provides a detailed description of the methods when using them for 2-channel sleep staging and brain computer interfacing tasks. Gaussian noise DA has been used in many studies [4], [5], [11], [18] and involves adding Gaussian noise to the training data. We used a mean of 0 and standard deviations (SD) between 0.1 and 1.0 in increments of 0.1 SD. Time reverse DA involves flipping the training samples over the time dimension such that the first point in a sample becomes its final point and vice versa. Smooth time masking involves zeroing out a window of signal within a sample, where the edges of the window are tapered using a pair of sigmoid functions. The masking has two parameters: a temperature parameter that controls the steepness of tapering, which we set to 0.5, and a length parameter that controls the length of the signal that is zeroed out for which we tried values between 0.2 and 2 seconds (s) in 0.2s increments. We applied identical perturbations to each channel. For FT surrogate DA, we randomized the phase of the samples by shifting the phase of each frequency by values selected randomly from a uniform distribution between 0 and a maximum phase perturbation. We examined 10 maximum phase perturbation linearly interspersed between 0 and 2π radians. For the frequency shift DA, we shifted each frequency by a value randomly selected from a uniform distribution between [-f_max_, +f_max_]. We examined 10 f_max_ values linearly distributed between 0 and 3Hz. Each sample was perturbed separately, and the same perturbation was applied to each of a sample’s respective channels. For channel dropout, we zeroed out channels selected via Bernoulli random variables in 10% increments between 10% and 90%.

### C. Model Development Approach

To determine whether DA improved model performance, we performed family-wise, paired, two-tailed t-tests between the performance of each model. We then applied separate false discovery rate (FDR) correction to the p-values of each metric.

### D. Spectral Explainability Analysis

Given that many of the DA approaches were frequency-based, we were curious how they may have affected the importance of each canonical frequency band. As such, we used a spectral explainability approach similar to that used in [6][7]. We obtained the model test ACC in each fold, applied a fast Fourier transform (FFT) to convert the test samples to the frequency domain, replaced frequency band-specific Fourier coefficients with values of zero, converted back to the time domain with an inverse FFT, obtained the model ACC on the perturbed samples, and calculated the percent change (PCT_CHG) in ACC. We used the δ (0-4 Hz), θ (4-8 Hz), α (8-12 Hz), β (12-25 Hz), γ_1_ (25-45 Hz), γ_2_ (55-75 Hz), and γ_3_ (75-100 Hz) bands. We divided the γ-band into 3 sub-bands, where γ_1_ and γ_2_ were divided due to line noise and γ_2_ and γ_3_ were divided to limit the effects of their perturbation to a smaller band. We applied this analysis to all folds and models.

### E. Spatial Explainability Analysis

Some of the DA methods may have affected the importance of individual channels, so we used a spatial explainability approach to examine this possibility [6]. We ablated (i.e., replaced with zeros) individual channels in the test set and obtained the PCT_CHG in ACC resulting from the ablation. Afterwards to identify the top channels across models, we counted the percentage of models for which each channel was among the top 10 channels (i.e., based on mean absolute importance across folds) and selected those 10 channels that most commonly occurred in the top 10 channels.

## III. Results and Discussion

In this section, we describe and discuss our model performance and explainability analysis results.

### A. Model Performance Analysis

Table 1 shows our model performance results. While there are some visually distinguishable differences in the means of model performance, we did not observe any statistically significant differences in performance after correction. That is likely attributable to both the large number of models to which we applied statistical testing and the relatively small number of performance values (i.e., one value per each of 25 folds).

**TABLE 1.** Model Performance Results.

| Model | ACC | BACC | SENS | SPEC |
| --- | --- | --- | --- | --- |
| M1.1 | 86.30±12.28 | 85.84±15.94 | 92.79±16.99 | 78.89±28.18 |
| M1.2 | 88.75±10.90 | 88.23±14.66 | 92.67±11.51 | 83.80±26.99 |
| M1.3 | 87.51±11.32 | 87.85±11.27 | 91.64±18.86 | 84.05±19.28 |
| M2.1 | 88.66±10.30 | 88.61±12.96 | 91.62±12.24 | 85.60±22.98 |
| M2.2 | 87.81±11.09 | 86.99±15.55 | 92.33±13.71 | 81.66±28.59 |
| M3 | 84.68±13.34 | 84.27±16.50 | 91.58±17.45 | 76.97±29.30 |
| M4.1 | 88.28±11.56 | 89.23±11.26 | 92.55±14.72 | 85.91±20.55 |
| M4.2 | 88.78±11.69 | 87.84±15.67 | 92.56±13.84 | 83.13±27.76 |
| M5.1 | 88.39±11.02 | 88.13±13.16 | 91.27±13.98 | 84.99±22.82 |
| M5.2 | 87.95±12.31 | 87.26±15.79 | 91.53±14.56 | 83.00±28.54 |
| M6.1 | 88.82±10.89 | 88.53±13.40 | 92.34±12.26 | 84.72±23.46 |
| M6.2 | 86.79±12.04 | 86.67±15.12 | **92.80±11.12** | 80.54±28.12 |
| M7.1 | 89.05±11.14 | 87.96±15.05 | 90.74±19.46 | 85.18±24.30 |
| M7.2 | **90.75±10.14** | **89.72±12.51** | 92.35±18.74 | **87.08±19.73** |

Based on visual inspection of mean model performance, with the exception of time reverse DA (M3), most DA methods improved performance relative to training on one copy of the training data (M1.1). Nevertheless, duplicating the training data (M1.2) led to comparable model performance, which suggests that future studies might benefit from simply duplicating their training data when training a model and that future studies developing EEG DA approaches should consider comparing the performance of models trained with their new approaches to models trained on duplicate training data.

Tripling dataset size (i.e., two augmented copies of data) seemed to mostly result in slightly decreased performance relative to just training with the un-augmented data and a single copy of augmented data. It is possible in this case that, even with different random initializations, training with two copies of data augmented in the same manner did not introduce enough variation into the data for the model to learn more generalizable patterns. Nevertheless, tripling dataset size with channel dropout augmentation (M7.2) did lead to improved model performance relative to the other DA approaches and other studies on the same dataset [6], [7].

### B. Spectral Explainability Analysis

Figure 2 shows our spectral explainability results across models. Note that γ_2_ and γ_3_ were relatively unimportant to model predictions, which is important given that the data was originally low pass filtered below 70 Hz. Across models, δ, α, and β tended to be highly important. There is some overlap between our results and those of [6], which identified δ, β, and γ importance, and those of [7], which identified δ, θ, and β importance. Spectral importance did not seem to change very much when datasets were doubled versus tripled, though β importance did seem to change notably between M5.1 and M5.2. The frequency domain augmentations in M5.1 seemed to reduce the impact of perturbing the β-band, though M5.2 and M6.1 and M6.2 did not have a similar effect. This is an interesting finding given that M5.1 did not seem to have a noticeable difference in performance relative to M5.2, M6.1, and M6.2 and would suggest that the FT Surrogate approach of M5.1 had the potential to increase model robustness.

**Figure 2.**
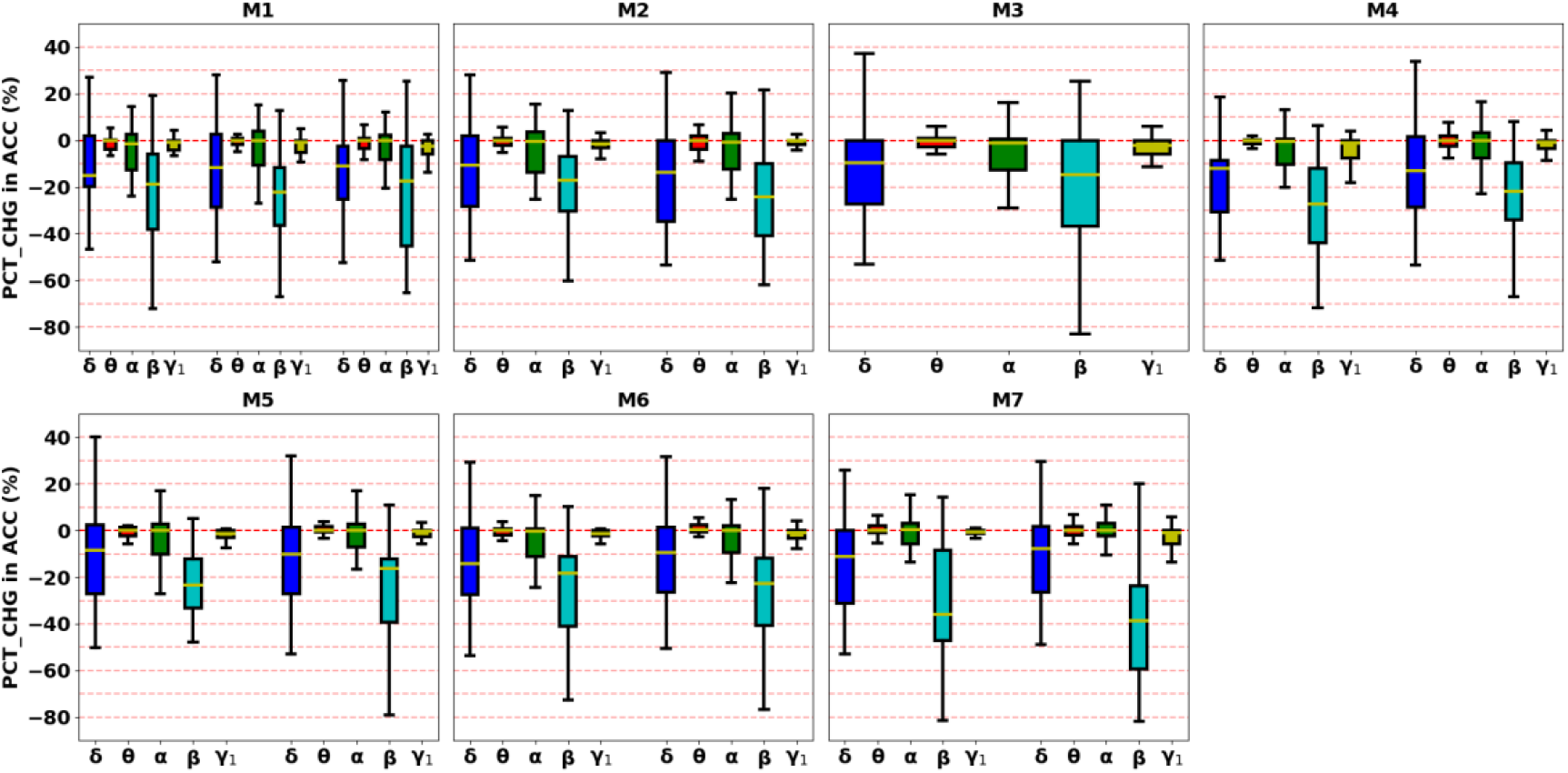
Spectral Explainability Results. Each panel shows results for models trained with a different DA approach. In each panel, importance values are further displayed from left to right in order of the number of data duplicates (i.e., double or tripled training set size). The frequency bands for each model are on the x-axis, and the PCT_CHG in ACC after perturbation is on the y-axis and aligned across panels.

### C. Spatial Explainability Analysis

Figure 3 shows our spatial explainability results and the location of each electrode. Interestingly, the majority of the top 10 electrodes that we identified were in the left hemisphere (F7, F3, T3, T5, P3) or were central electrodes (Fz, Cz, Pz), though we did identify two right hemisphere electrodes (F4, P4). Previous studies have found that reduced left hemisphere activity in MDD [19]. Interestingly, copying the training data multiple times (M1.1-1.3) tended to increase the effects of channel loss, where the model perhaps learned to rely more upon each EEG channel. In contrast, increasing the number of augmented copies of data from 1 to 2 copies did not seem to lead to a similar increased effect of channel perturbation (M2-M7). Among the DA approaches that affected the time and spatial dimensions of the data, channel dropout DA reduced the impact of channel perturbation to negligible levels, and time reverse and smooth time masking DA did not seem to impact model sensitivity to channel perturbation.

**Figure 3.**
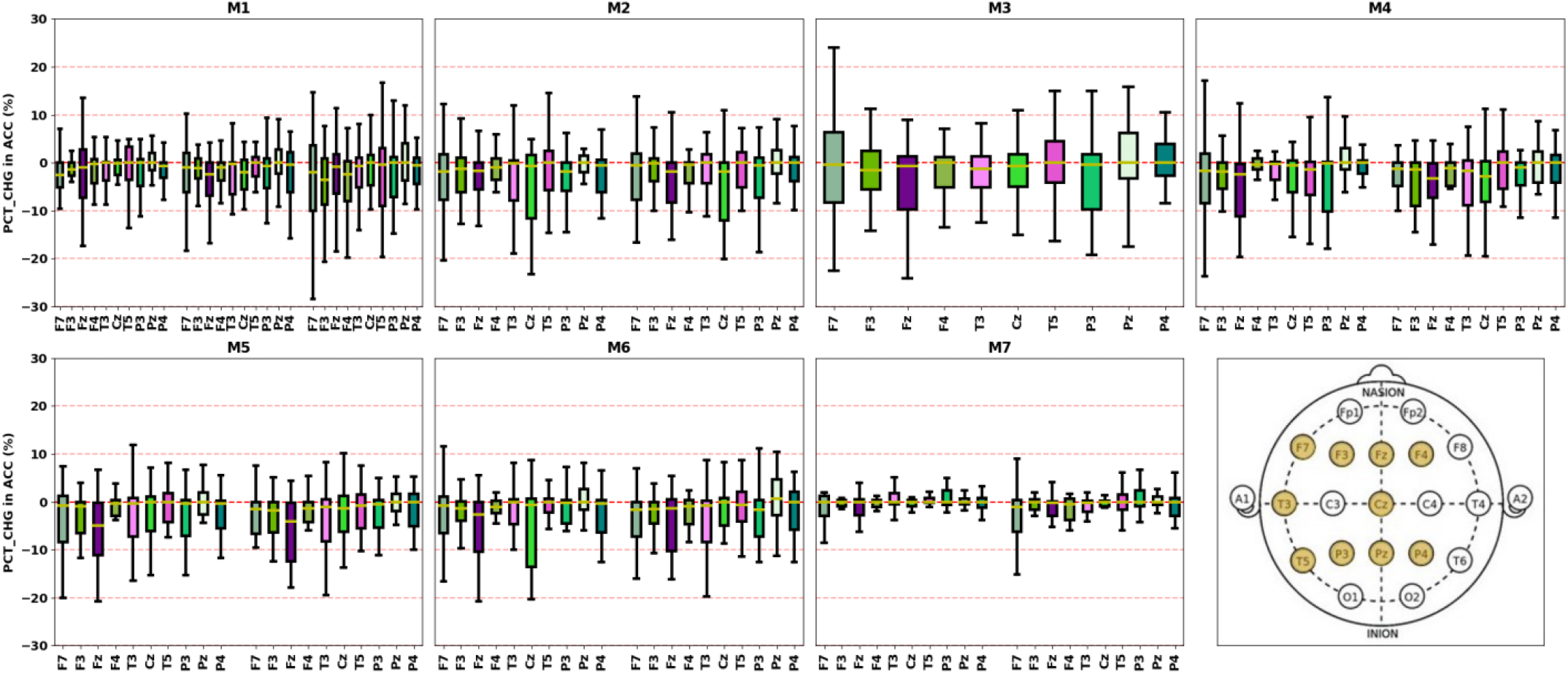
Spatial Explainability Results. Each panel shows results for models trained with a different DA approach. In each panel, importance values are further displayed from left to right in order of the number of data duplicates (i.e., double or tripled training set size). The top 10 channels are on the x-axis, and the PCT_CHG in ACC after perturbation is on the y-axis and shared across panels. The bottom right panel shows the location of each channel on the scalp, with the top 10 channels highlighted.

### D. Limitations and Future Work

We tested all DA approaches independently. However, training with different combinations of DA approaches and DA parameters could prove helpful. Importantly, our findings on the utility of each DA approach are restricted to our dataset and model. Future studies investigating the relative utility of DA techniques should apply them to multiple datasets and architectures. Moreover, other EEG DA approaches have been developed in previous studies, and applying them to our dataset and classifier could be helpful.

## IV. Conclusion

Deep learning methods are increasingly being applied to EEG data for neuropsychiatric disorder diagnosis in research settings. However, the development of reliable models needed for clinical translation is hindered by small dataset size. EEG DA approaches could address this concern. In this study, we explore the use of 6 different EEG augmentation approaches for improving MDD diagnosis performance and also compare the utility of the DA approaches relative to a new baseline of training on duplicate training data. We further examine the effects that they have upon the representations learned by the models through a series of explainability analyses. We find that most approaches improve performance but not relative to simply duplicating the training set. The exception to this is channel dropout DA, which does improve performance. These findings suggest that future studies developing EEG DA methods would benefit from comparing to a baseline of duplicate training data to account for the discrepancy in training set size and determine whether useful augmentations are introduced. While restricted to our dataset and model architecture, we hope that our findings will guide future deep learning studies on small EEG datasets and guide the further development of EEG augmentation approaches.

## References

[1] C. Rommel, J. Paillard, T. Moreau, and A. Gramfort, “Data augmentation for learning predictive models on EEG: a systematic comparison,” J. Neural Eng., vol. 19, no. 6, 2022, doi: 10.1088/1741-2552/aca220.

[2] Z. Wan, R. Yang, M. Huang, N. Zeng, and X. Liu, “A review on transfer learning in EEG signal analysis,” Neurocomputing, vol. 421, pp. 1–14, 2021, doi: 10.1016/j.neucom.2020.09.017.

[3] X. Zhou et al., “Interpretable and Robust AI in EEG Systems: A Survey,” pp. 1–18, 2018.

[4] C. A. Ellis, A. Sattiraju, R. L. Miller, and V. D. Calhoun, “A Framework for Systematically Evaluating the Representations Learned by A Deep Learning Classifier from Raw Multi-Channel Electroencephalogram Data,” bioRxiv, 2023.

[5] C. A. Ellis, A. Sattiraju, R. L. Miller, and V. D. Calhoun, “Novel Approach Explains Spatio-Spectral Interactions in Raw Electroencephalogram Deep Learning Classifiers,” IEEE International Conference on Acoustics, Speech, and Signal Processing Workshops, 2023.

[6] C. A. Ellis, R. L. Miller, and V. D. Calhoun, “Improving Multichannel Raw Electroencephalography-based Diagnosis of Major Depressive Disorder via Transfer Learning with Single Channel Sleep Stage Data,” bioRxiv, 2023.

[7] C. A. Ellis, R. L. Miller, and V. D. Calhoun, “Cross-Sampling Rate Transfer Learning for Enhanced Raw EEG Deep Learning Classifier Performance in Major Depressive Disorder Diagnosis,” in bioRxiv, 2023, pp. 2–6.

[8] K. Simonyan and A. Zisserman, “Very Deep Convolutional Networks for Large-Scale Image Recognition,” in International Conference on Learning Representations (ICLR), 2015, pp. 1–14.

[9] W. Mumtaz, L. Xia, M. A. M. Yasin, S. S. A. Ali, and A. S. Malik, “A wavelet-based technique to predict treatment outcome for Major Depressive Disorder,” PLoS One, vol. 12, no. 2, pp. 1–30, 2017, doi: 10.1371/journal.pone.0171409.

[10] H. W. Loh, C. P. Ooi, E. Aydemir, T. Tuncer, S. Dogan, and U. R. Acharya, “Decision support system for major depression detection using spectrogram and convolution neural network with EEG signals,” Expert Syst., vol. 39, no. 3, pp. 1–15, 2022, doi: 10.1111/exsy.12773.

[11] C. A. Ellis, A. Sattiraju, R. Miller, and V. Calhoun, “Examining Effects of Schizophrenia on EEG with Explainable Deep Learning Models,” in 2022 IEEE 22nd International Conference on Bioinformatics and Bioengineering (BIBE), 2022, pp. 301–304. doi: 10.1109/BIBE55377.2022.00068.

[12] S. L. Oh, J. Vicnesh, E. J. Ciaccio, R. Yuvaraj, and U. R. Acharya, “Deep convolutional neural network model for automated diagnosis of Schizophrenia using EEG signals,” Appl. Sci., vol. 9, no. 14, 2019, doi: 10.3390/app9142870.

[13] T. O’Malley, E. Bursztein, J. Long, F. Chollet, H. Jin, and L. Invernizzi, “KerasTuner,” 2019.

[14] C. Rommel, T. Moreau, J. Paillard, A. Gramfort, and U. Paris-saclay, “CADDA : Class-wise Automatic Differentiable Data Augmentation for EEG Signals,” 2022.

[15] M. N. Mohsenvand, P. Maes, E. E. Alsentzer, M. B. A. Mcdermott, F. Falck, and S. K. Sarkar, “Contrastive Representation Learning for Electroencephalogram Classification,” Proc. Mach. Learn. Res., vol. 136, pp. 238–253, 2020.

[16] J. T. C. Schwabedal, J. C. Snyder, A. Cakmak, S. Nemati, G. D. Clifford, and S. P. Jan, “Addressing Class Imbalance in Classification Problems of Noisy Signals by using Fourier Transform Surrogates,” pp. 1–8, 2019.

[17] A. Saeed, D. Grangier, O. Pietquin, and N. Zeghidour, “Learning from heterogeneous EEG signals with differentiable channel reordering,” in IEEE International Conference on Acoustics, Speech and Signal Processing - Proceedings, 2021, pp. 1255–1259. doi: 10.1109/ICASSP39728.2021.9413712.

[18] A. Sattiraju, C. A. Ellis, R. L. Miller, and V. D. Calhoun, “An Explainable and Robust Deep Learning Approach for Automated Electroencephalography-based Schizophrenia Diagnosis,” 2023.

[19] D. Hecht, “Depression and the hyperactive right-hemisphere,” Neurosci. Res., vol. 68, no. 2, pp. 77–87, 2010, doi: 10.1016/j.neures.2010.06.013.

